# A hydrophobic network: Intersubunit and intercapsomer interactions stabilizing the bacteriophage P22 capsid

**DOI:** 10.1101/626804

**Authors:** Kunica Asija, Carolyn M. Teschke

## Abstract

dsDNA tailed phages and herpesviruses assemble their capsids using coat proteins that have the ubiquitous HK97 fold. Though this fold is common, we do not have a thorough understanding of the different ways viruses adapt it to maintain stability in various environments. The HK97-fold E-loop, which connects adjacent subunits at the outer periphery of capsomers, has been implicated in capsid stability. Here we show that in bacteriophage P22, residue W61 at the tip of the E-loop plays a role in stabilizing procapsids and in maturation. We hypothesize that a hydrophobic pocket is formed by residues I366 and W410 in the P-domain of a neighboring subunit within a capsomer, into which W61 fits like a peg. In addition, W61 likely bridges to residues A91 and L401 in P-domain loops of an adjacent capsomer, thereby linking the entire capsid together with a network of hydrophobic interactions. There is conservation of this hydrophobic network in the distantly related P22-like phages, indicating that this structural feature is likely important for stabilizing this family of phages. Thus, our data shed light on one of the varied elegant mechanisms used in nature to consistently build stable viral genome containers through subtle adaptation of the HK97 fold.

**IMPORTANCE:** Similarities in assembly reactions and coat protein structures of the dsDNA tailed phages and herpesviruses make phages ideal models to understand capsid assembly and identify potential targets for antiviral drug discovery. The coat protein E-loops of these viruses are involved in both intra-and intercapsomer interactions. In phage P22, hydrophobic interactions peg the coat protein subunits together within a capsomer, where the E-loop hydrophobic residue W61 of one subunit packs into a pocket of hydrophobic residues I366 and W410 of the adjacent subunit. W61 also makes hydrophobic interactions with A91 and L401 of a subunit in an adjacent capsomer. We show these intra-and intercapsomer hydrophobic interactions form a network crucial to capsid stability and proper assembly.

## INTRODUCTION

There are several similarities in the assembly pathways of tailed bacteriophages and herpesviruses. *In vivo*, the assembly process is likely initiated with the dodecameric portal complex and involves the co-polymerization of multiple copies of the major capsid protein with a protein assembly catalyst, scaffolding protein (1). Following the formation of the DNA-free precursor capsid (procapsid), the DNA is packaged through the portal ring resulting in increases in the capsid volume and stability. This step is generally termed capsid expansion or maturation (2). The addition of the tail machinery marks the end of assembly (2). The three-dimensional arrangement of coat proteins in dsDNA viral capsids results in robust structures that can withstand high levels of internal pressure that is exerted by packaged DNA, which results in repulsive forces as well as the energetic strain imposed from DNA bending (3–5). In phages such as phi29 the pressure inside a capsid is known to exceed 6 MPa (6). In some phages, this pressure is thought to be necessary to eject the DNA into the prokaryotic host.

The presence of a common Hong Kong 97 fold (HK97 fold) in the coat proteins of tailed dsDNA phages and herpesviruses allows them to be grouped in a taxonomic lineage known as the HK97-like viruses (7). The lower domain of the Herpes Simplex virus 1 (HSV-1) coat protein also adopts the HK97 fold (8). This motif is the fundamental structure of these viral coat proteins, but also sometimes has the addition of an inserted domain (I-domain) such as in phages P22, T7, and phi29 (8–14). During assembly, capsid proteins are programmed by scaffolding protein to form hexons (with 6 coat protein subunits) or pentons (with five coat protein subunits). The HK97 fold is characterized by the presence of an axial A-domain that protrudes into the center of the hexons and pentons providing intracapsomer stability, a peripheral P-domain involved in intercapsomer interactions, and an elongated E-loop, which can interact within a subunit, in addition to intra- and intercapsomer interactions (15, 16). This fold exhibits enormous versatility by the addition of domains and extensions (8–10, 15, 17–20).

The interactions between and within capsomers during assembly allow for the formation of correctly sized procapsids and these initial interactions are often rearranged during the maturation reaction (16). Based on structural analyses of the HSV coat protein, multiple stabilizing interactions are made between the E-loop of the lower domain with other regions in the lower domain of the adjacent subunit (21). In phage HK97 virions, during maturation the E-loop of one capsid subunit is covalently crosslinked to the P-domain of an adjacent subunit, building concatemers of capsomers resulting in a stable capsid (15). Other non-covalent E-loop interactions are also crucial in assembly of HK97, including an E-loop to G-loop interaction across the icosahedral two-fold axes of symmetry. This interaction occurs only in procapsids and is broken during maturation. Phage T7 is proposed to stabilize capsomers through electrostatic interactions between its negatively charged E-loop and the positively charged surface of the P-domain of an adjacent subunit in the capsomer (22). T4 bacteriophage employs electrostatic forces both within and between capsomers to stabilize the capsid (23).

The P22 coat protein has the HK97 fold with an I-domain inserted into the A-domain (13, 24, 25) (Figure 1A). The D-loops in the I-domains make intercapsomer contacts across the 2-fold axis of symmetry to stabilize procapsids (26). Conformational changes in the P-loop, P-domain and the N-arm of the coat protein of P22 during the maturation reaction have also been proposed to play a role in stabilizing the capsid (13). Additional conformational changes occur during maturation at the level of capsomers. Prior to maturation, procapsid hexons are skewed and the coat proteins are compacted such that the walls of the procapsids are thicker than in virions (16, 25, 27). During maturation, the hexons rearrange so they become symmetric and flattened, leading to thinner walls in the virion. In addition, many inter-subunit contacts along the periphery of the A-domain must be broken and new ones formed during maturation (28, 29), so other capsomer contacts must remain intact for the particles to survive maturation.

**Figure 1.**
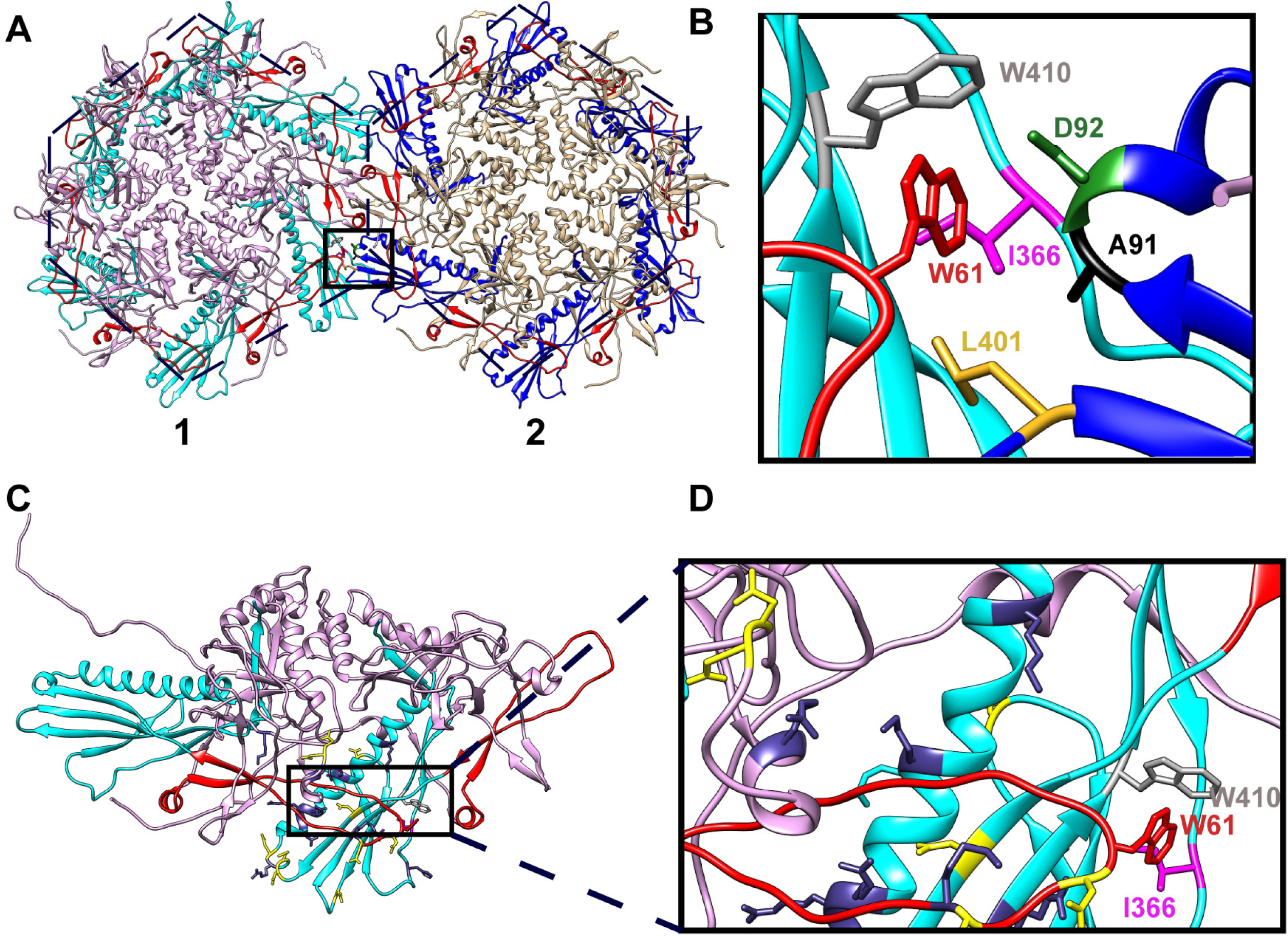
A hydrophobic peg to stabilize the P22 capsid. **A.** Neighboring capsomers in mauve and tan depicted by the numbers 1 and 2, respectively. The E-loops have been highlighted in red and the P-domains have been highlighted in cyan for capsomer 1 and blue for capsomer 2. The box shows the interacting residues forming the hydrophobic peg and pocket. **B.** Zoomed in inset showing W61 (red) from one subunit interacting with W410 (gray) and I366 (magenta) in the P-domains of the adjacent subunit (within capsomer 1). W61 also potentially interacts with A91 (black), D92 (green) and L401 (yellow), which are in the P-domains of the adjacent subunit. **C.** Neighboring coat protein subunits shown within capsomer 1. Inset shows the interacting residues from E-loop of one subunit and P-domain of adjacent subunit, within the capsomer. **D.** W61 (red) in E-loop of one subunit interacts with W410 (gray) and I366 (fuchsia) in P-domain of adjacent subunit. This interaction is reinforced by a cage of charged residues. Positively charged residues are shown in dark blue while negatively charged residues are shown in yellow.

As indicated above, the E-loop is suggested to universally make intra-capsomer connections to the P-domain of an adjacent subunit. Despite suggestions based on the recent 3.3 Å cryoEM structure of P22’s capsid (30), salt bridges formed between E-loop and P-domain of neighboring subunits are not important for assembly of PC or maturation into virions (Asija and Teschke, submitted). Here we investigate how the E-loop is used in the assembly of P22 procapsids and virions, focusing on the role of W61 located at the very tip of the E-loop. W61 is shown to make several intracapsomer hydrophobic contacts. Our data suggest a ring of charged residues surrounding this hydrophobic patch might strengthen this intra-capsomer interaction. We further show that W61 likely makes contacts with residues in the P-domain of an adjacent capsomer across two-fold axes of symmetry. Our data are consistent with the idea of universal, yet adaptable, stabilizing contacts made by the E-loop of the HK97 fold.

## RESULTS

### Residue W61 in the E-loop plays a role in capsid stabilization and maturation

As we have shown previously, several salt bridges formed between E-loop and P-domain are not crucial for capsid integrity (Asija and Teschke, submitted). However, we noted the presence of W61 at the tip of the E-loop and hypothesized that it could be making stabilizing hydrophobic interactions with the P-domains of an adjacent subunit within a capsomer, as well as between capsomers (Figure 1). Tryptophans have been suggested to be important in loops that are involved in protein:protein interfaces (31). Thus, we assessed the role of W61 in capsid assembly and stability by mutagenesis of the site, characterization of subsequent phenotype, and stability of the capsids.

#### All the W61 substitutions except W61Y cause a lethal phenotype

The effects of polar, non-polar and aromatic amino acid substitutions at this site were determined. W61Y, W61N and W61V substitutions were generated using site-directed mutagenesis of plasmid-encoded gene 5, which codes for coat protein. We complemented a 5^*-*^*am* phage with expression of the WT and W61 variant coat proteins from the plasmid at different temperatures to determine if the substitutions caused any phenotype by efficiency of plate (EOP), as described in the Materials and Methods. In this assay, plaques are only formed when the expressed coat protein variant can be assembled into viable phages at the tested temperatures (Figure 2A, Table 2). Coat protein with the W61Y substitution was able to generate phages at all temperatures, similar to that of WT coat protein. Conversely, complementation by coat proteins carrying substitutions W61N and W61V resulted in a lethal phenotype at all temperatures, as plaques were found only near the reversion frequency of the *5*^*-*^*am* phage stock. These data show that residue W61 is important *in vivo* phage P22 production.

**Figure 2.**
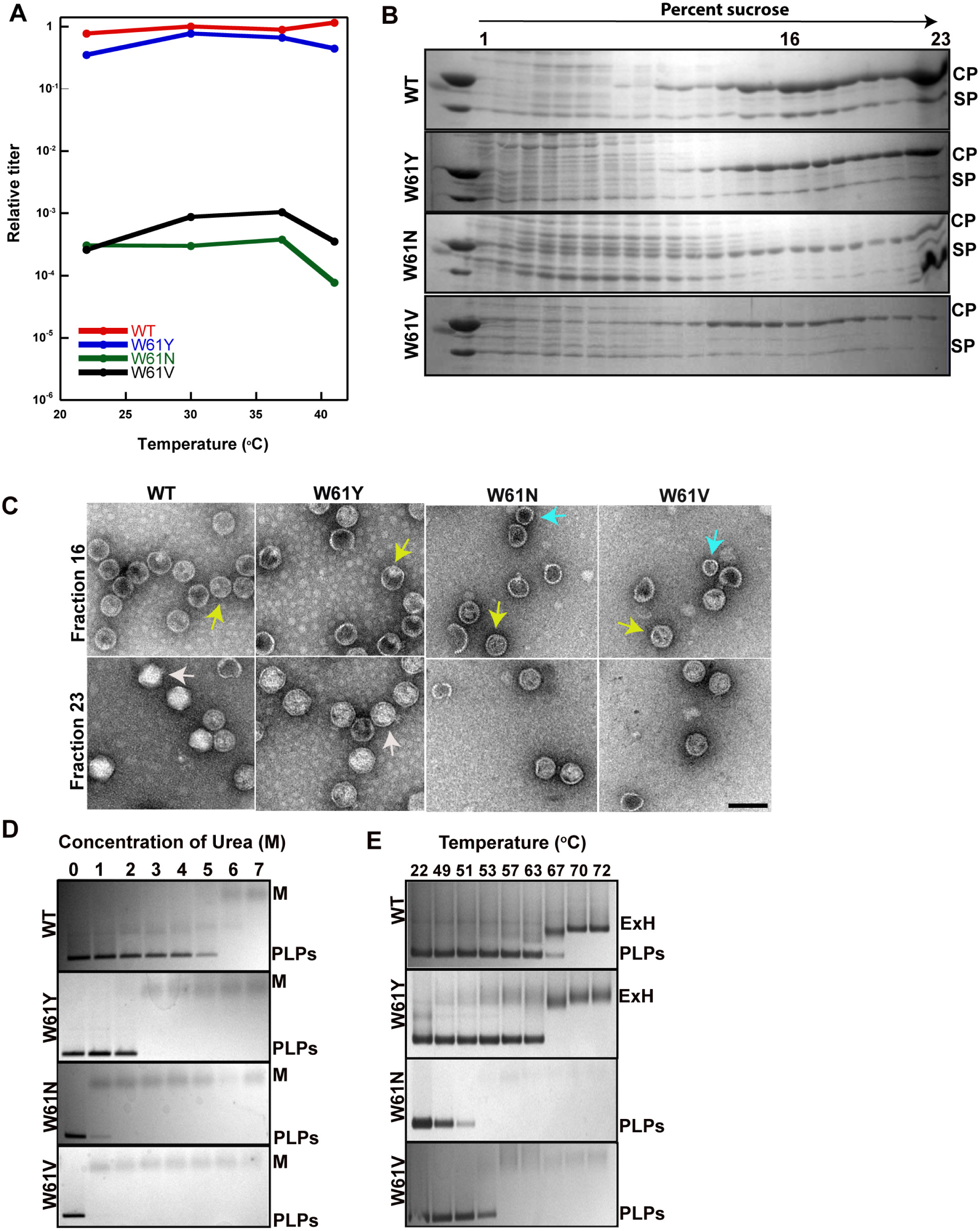
W61 in the E-loop stabilizes the P22 capsid. **A.** Titers of phages assembled by complementation with W61Y (blue), W61N (green) and W61V (black) coat protein substitutions relative to those with WT (red) coat protein at 30 °C. **B.** SDS gels of 5-20% sucrose density gradient fractions from a phage-infected cell lysate to separate procapsids and phages with WT coat protein and coat protein with the W61 substitutions. Fraction 16 is where normal procapsids sediment and fraction 23 is where phages and other particles are found. CP, coat protein; SP, scaffolding protein. **C. Top row**. TEM images of procapsid fractions (fraction 16) from sucrose density gradient of WT and W61 coat variants. **Bottom row**. Mature phage fractions (fraction 23) from the sucrose density gradient observed under the TEM. Yellow arrows show T=7 sized procapsids, cyan arrow show petite procapsids and white arrows are highlighting mature phages. Scale bar is 100 nm. **D.** Stabilities of PLPs assembled with WT, W61Y, W61N and W61V coat proteins tested by treating with varying concentrations of urea (0-7M). Samples were run on a 1% native agarose gel. PLPs, Procapsid-like particles; M, unfolded monomers. **E.** PLPs with WT, W61Y, W61N and W61V coat proteins were subjected to temperatures ranging from 22 °C to 72 °C to induce heat dependent expansion. Samples were run on a 1% agarose gel. ExH, heat-expanded procapsids.

#### Lethal effect of W61 substitutions is due to their inability to mature into infectious virions

To test whether the lethal effect of the W61N and W61V substitutions is caused by an assembly defect or a maturation defect, we isolated particles from phage-infected cells. Procapsids and mature virions, produced from a 5^-^13^-^*am* infection complemented by expression of WT or mutant coat protein from a plasmid, were separated by a 5-20% sucrose density gradient. In Figure 2B, 10% SDS gels of the sucrose gradient fractions are shown. WT and coat proteins with W61 substitutions generate procapsids (peak centered at about fraction 16) composed both of coat protein and scaffolding protein. The procapsids also contain the ejection proteins and the portal protein complex (data not shown). However, the procapsids of coat protein variants W61N do not have the usual amount of scaffolding protein. The procapsid peak for the lethal substitutions (W61N and W61V) was shifted up the gradient to lower sucrose concentrations, suggesting the mass of the particles had decreased. This is usually due to decreased content of scaffolding proteins, particles of smaller diameter or incomplete particles (32–34). Fractions 16 and 23 were observed using negative stain transmission electron microscopy (TEM) and showed normal sized procapsids (yellow arrows) in fraction 16 and mature virions (white arrows) in fraction 23 for WT and W61Y coat proteins. (Figure 2C). Coat proteins with lethal substitutions,W61N and W61V formed normal sized procapsids (yellow arrows), as well as petite procapsids (cyan arrows), but these were unable to mature into infectious virions (Figure 2C).

#### Stability of procapsid-like particles with W61N and W61V substitutions is drastically decreased

Since W61N and W61V substitutions confer lethal phenotypes, we hypothesized the substitutions could be destabilizing the procapsids considering that the E-loop is usually involved in both intra-and intercapsomer interactions. To test this, we determined the urea concentration required to dissociate procapsid-like particles (PLPs, composed of solely of coat and scaffolding proteins expressed from a plasmid), which also results in the unfolding of coat and scaffolding proteins as described in the Methods. PLPs assembled from WT coat protein denatured to monomers at 5-6 M urea (Figure 2D). W61Y PLPs are less stable, unfolding at 3 M urea, than WT PLPs despite having a WT-like phenotype. Lethal W61N and W61V coat protein variant PLPs are very destabilized, denaturing at concentrations as low as 1 M urea. These data show that size, and perhaps the hydrophobicity, of the amino acid is important at this position in order to maintain stabilizing interactions with the other residues.

#### Residue W61 is crucial to capsid expansion

To determine if the W61 variant PLPs can undergo maturation, we used an *in vitro* heat expansion assay, as described in Methods. When PLPs expand, there is an increase in the diameter, leading to slower migration of the particles on an agarose gel (35). The PLPs assembled with WT coat protein or the non-lethal W61Y coat protein mutant heat expanded at ∼ 67 °C (Figure 2E). The lethal W61N and W61V variants are unable to undergo normal capsid maturation and instead rupture at 53-57 °C. In total, these data indicate that the E-loop residue W61 makes important contacts for PLP stability and ability to expand.

#### There is no effect on secondary structure of W61 coat variants and their interaction with scaffolding protein

Our data above (Figure 2B) showed that some PCs assembled from W61 coat protein variants had less than normal scaffolding protein content, which is somewhat puzzling as the E-loop is on the exterior of PCs. Two possibilities seem reasonable. 1) The coat protein monomer conformation could be affected by the amino acid substitutions at position 61 such that there was decreased interaction with scaffolding protein during assembly, or 2) the assembled coat protein shell had decreased affinity for scaffolding protein. We tested the interaction between coat protein monomers and scaffolding protein using weak affinity chromatography, where scaffolding protein is attached to a column matrix and coat protein is passed over this matrix. The monomeric coat protein is retained compared to a control protein (ovalbumin) when it interacts with scaffolding protein on the column (36, 37). WT coat monomers, as well as all of the W61 coat variants, interacted with scaffolding protein as seen by their retention on the column as compared to when non-interacting ovalbumin is applied (Figure 3A). We determined there was no significant change in secondary structure for any W61 variant coat protein monomers as compared to WT coat protein by circular dichroism spectroscopy (Figure 3B). Finally, we tested the ability of each coat protein to assemble *in vitro* by adding scaffolding protein to coat protein monomers (Figure 3C). Each of the W61 variant coat proteins was able to assemble *in vitro*, albeit with decreased efficiency compared to WT coat protein. Thus, the amino acid substitutions at W61 appear to affect the structure of procapsids, rather than monomers.

**Figure 3.**
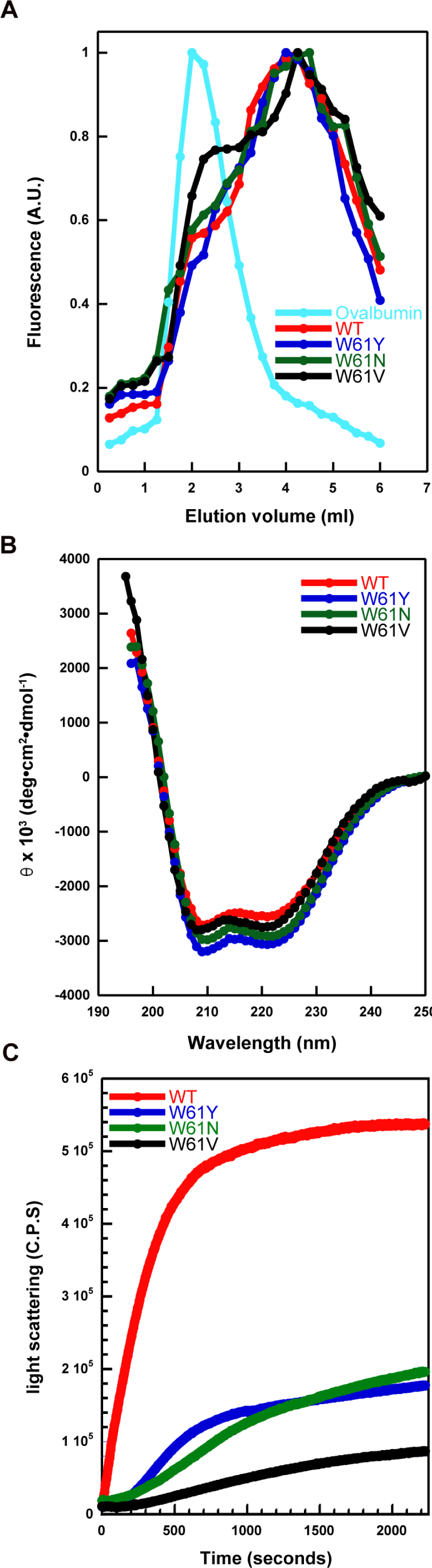
W61 coat substitutions do not affect the coat protein secondary structure or interaction with scaffolding protein. **A.** Weak affinity chromatography. The volume at which the WT coat monomer or with W61Y (blue), W61N (green), W61V (black) eluted (elution volume) was determine by tryptophan fluorescence recorded in arbitrary units (A.U). The monomers were run over a cobalt IMAC column charged with his-tagged scaffolding protein. Ovalbumin (cyan) was used as a negative control. **B.** Circular dichroism. Secondary structure analysis using circular dichroism of coat monomer with WT sequence (red) and W61 coat variants with W61Y, W61N and W61V shown in blue, green and black respectively. **C.** *In vitro* assembly of PLPs. Light scattering of WT coat monomers (red) along with the other coat variants was tested by incubating with scaffolding protein.

### Hydrophobic interactions between the E-loop and the P-domain of the adjacent subunit within a capsomer are crucial for capsid stability

We used the 3.3 Å cryoEM structure (PDB file 5UU5) (30) to determine which amino acids residue W61 could be interacting with in the adjacent subunit of a capsomer. Based on this structural assessment, we hypothesize that residues I366 (4.87 Å from W61) and W410 (4.34 Å from W61) in the adjacent subunit could be forming a hydrophobic pocket into which W61 inserts to stabilize the particles (Table 1, Figure 1). Thus, we made substitutions at these sites to test our hypothesis.

**Table 1.** Predicted distances between amino acid pairs between subunits of the P22 capsomer.

| Amino acid pair | Nature of interaction | Distance (Å)<sup>a</sup> |
| --- | --- | --- |
| W61-I366 | Intracapsomer | 4.34 |
| W61-W410 | Intracapsomer | 4.87 |
| W61-A91 | Intercapsomer | 4.88 |
| W61-D92 | Intercapsomer | 3.68 |
| W61-L401 | Intercapsomer | 4.16 |

<sup>a</sup>Distance measurements were made using the software Chimera (38).

**Table 2.** Summary of mutants generated in this study, results based on their phenotypes and capsid stabilities as tested by urea titrations.

| **Substitution** | **Location on coat protein<sup>a</sup>** | **Phenotype** | **Concentration of urea (M) at which virions decline to 50% of original population** |
| --- | --- | --- | --- |
| WT coat protein | N/A | N/A | 3-3.5 |
| W61Y | E-loop | WT-like | 3.5 |
| W61N | E-loop | Lethal | N/A |
| W61V | E-loop | Lethal | N/A |
| I366R | P-domain | Lethal | N/A |
| I366A | P-domain | WT-like | 1.6 |
| I366D | P-domain | Lethal | N/A |
| W410R | P-domain | *cs* and *ts* | 1.1 |
| W410A | P-domain | WT | 1.5 |
| W410D | P-domain | *ts* | 3.9 |
| A91D | P-domain | *cs* | 1 |
| A91V | P-domain | WT-like | 3.4 |
| D92A | P-domain | WT-like | 3.1 |
| D92R | P-domain | WT-like | 3.2 |
| L401A | P-domain | *cs* and *ts* | 2.2 |
| L401D | P-domain | Lethal | N/A |

<sup>a</sup>Refer to Table 1 for the distance between potential interacting residues. N/A, not applicable.

#### Substitutions in the hydrophobic pocket lead to phenotypic changes

Site-directed mutagenesis was used to generate arginine, alanine and aspartic acid substitutions at residues I366 and W410 in plasmid-encoded gene 5 to test the effects of charged residues in addition to a non-polar amino acid substitution. The phenotype due to the expression of these coat protein variants in cells infected with a gene 5^-^*am* phage was determined, as described above, and the EOP. The I366A coat protein substitution resulted in essentially a WT phenotype, with a small drop in the relative titer at 22 °C (Figure 4A, Table 2). I366R and I366D coat protein variants caused a lethal phenotype, where the titers were observed only at the reversion frequency of the 5^-^*am* phage at all the temperatures tested.

**Figure 4.**
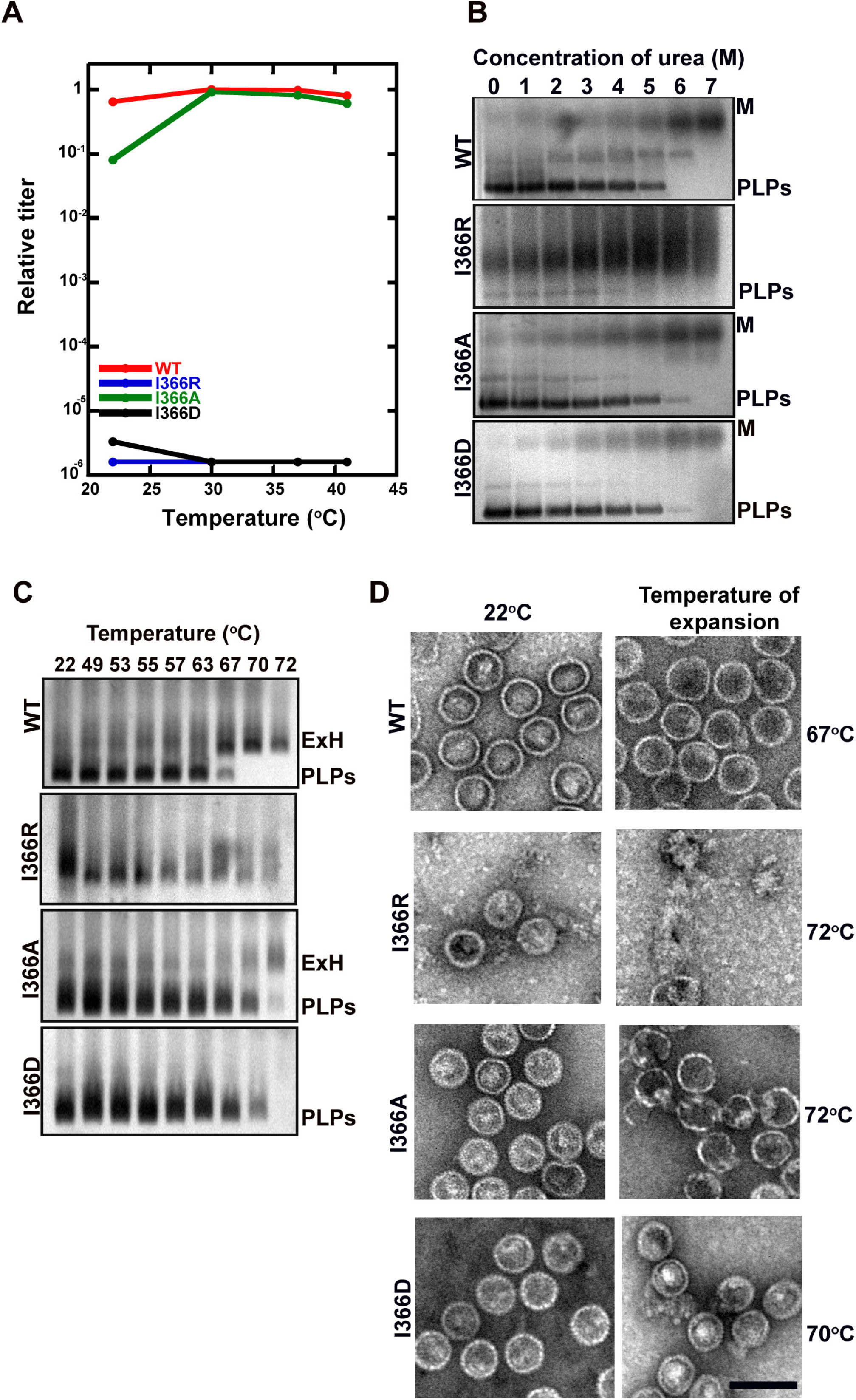
Charged substitutions at I366 result in a lethal phenotype. **A.** Relative titers of phages assembled by complementation with I366R (blue), I366A (green) and I366D (black) coat protein substitutions relative to phages with WT coat substitutions (red) at 30°C. **B.** Stabilities of PLPs made with WT, I366R, I366A and I366D coat proteins were tested by treating with varying concentrations of urea (0-7M). Samples were run on a 1% agarose gel. **C.** PLPs with WT, I366R, I366A and I366D coat proteins were subjected to temperatures ranging from 22 °C to 72 °C. Samples were then run on a 1% agarose gel to test for maturation. **D.** *In vitro* heat expanded PLPs with WT coat protein and coat protein with I366 substitution were observed under the electron microscope. The samples from the 22°C sample as well as the temperature at which they expand were observed under the microscope. Scale bar represents 100 nm. ExH, heat-expanded heads; M, unfolded monomers; PLPs, procapsid-like particles.

Coat proteins with substitutions at W410 cause less dramatic phenotypes. Coat protein W410A had a WT phenotype at all the temperatures tested (Figure 5A, Table 2). Coat protein variant W410R caused a decrease in titer at 22 °C, and therefore had a cold-sensitive (*cs*) phenotype, as well as a decrease in titer at 41 °C indicating the substitution caused a temperature-sensitive (*ts*) phenotype. The W410D variant showed a *ts* phenotype in the complementation assay. Thus, we can conclude that both residues I366 and W410 are important in phage production, and that this site is sensitive to a change of the residues to charged amino acids but the size of a non-polar residue is not critical.

**Figure 5.**
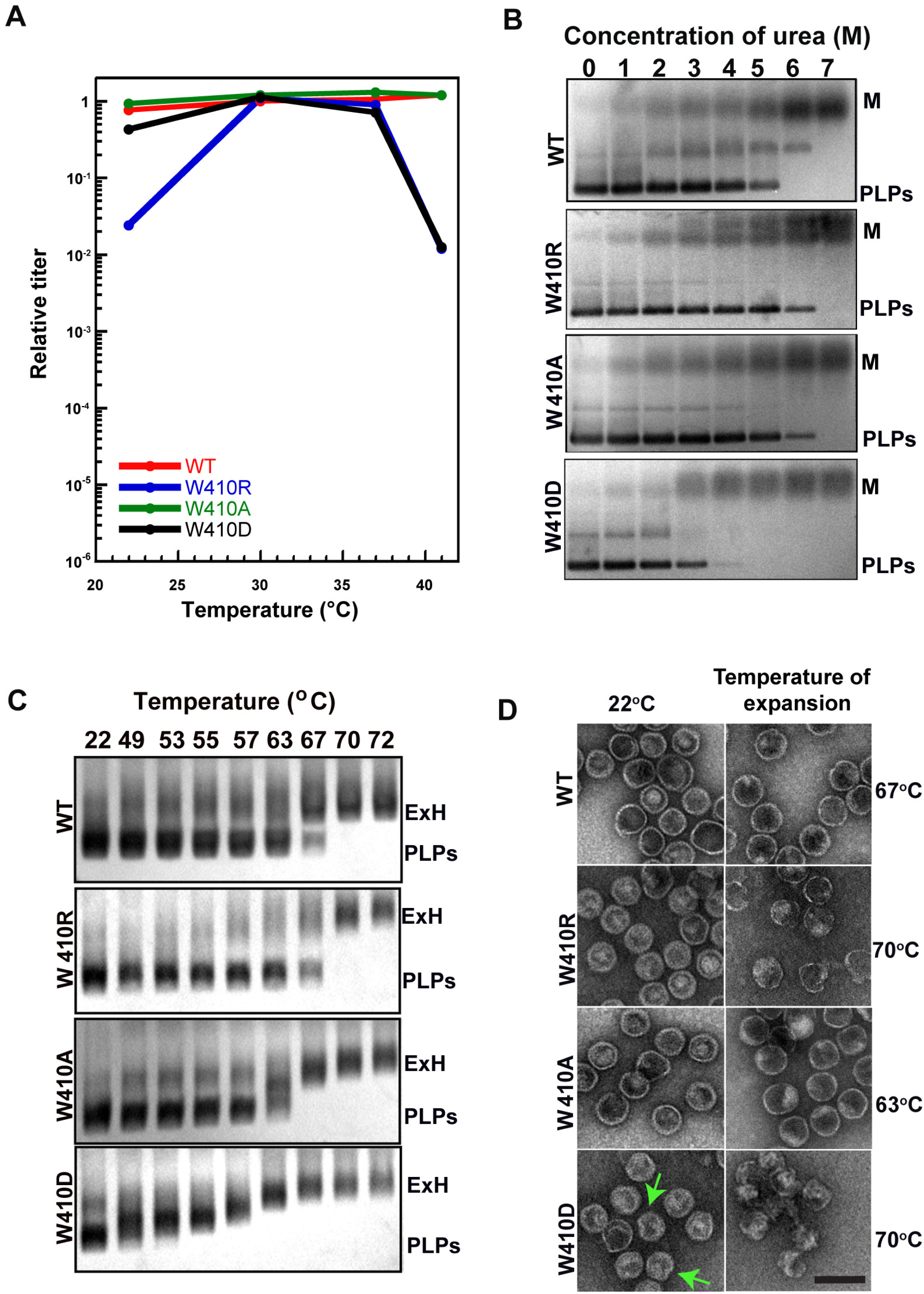
Charged substitutions of W410 cause phenotypes. **A.** Titers of phages assembled by complementation with coat protein W410R (blue), W410A (green), W401D (black) coat substitutions relative to WT coat protein (red) at 30°C. **B.** Stabilities of PLPs with WT coat protein and coat protein with W410R, W410A and W410D substitutions were tested by treating with varying concentrations of urea (0-7M). Samples were run on a 1% agarose gel. **C.** PLPs made with WT, W410R, W410 and W410D coat variants were subjected to temperatures ranging from 22 °C to 72 °C. Samples were then run on a 1% agarose gel to test for maturation. ExH, heat-expanded heads; PLPs, procapsid-like particles; M, unfolded monomers. D. Electron micrographs of *in vitro* heat-expanded PLPs with W410 coat substitutions. PLPs with substitutions were observed at 22°C and at the temperature at which they expanded/ruptured. Green arrows highlight aberrant particles formed at 22°C with the W410D coat protein substitution. Scale bar is 100nm.

#### The coat protein residues I366 and W410 are important for stability and capsid expansion of PLPs

A change in the stability of procapsids assembled with the I336 and W410 coat protein variants could explain the observed effects on phage biogenesis. Therefore, PLPs assembled from WT coat protein, coat protein with the I366A, R, D or W410A, R or D substitutions were tested for their stability to urea, as described above (Figures 4B, 5B). The temperature at which the PLPs expand was also determined.

PLPs with I366A and I366D coat substitutions denatured to form monomers at a urea concentration of 6 M, similar to WT PLPs (Figure 4B). In contrast, the I366R substitution appears to be an extremely destabilizing substitution. The yield of correctly sized PLPs was low and they denatured at 4-5 M urea (Figure 4B). There was also a substantial amount of smeared protein in the agarose gel from broken particles and aggregated coat protein. This was confirmed by TEM where the sample is composed mostly of aggregates and some PLPs (Figure 4D). PLPs made with W410R and W410A coat proteins denatured to monomers at 7 M urea, so were slightly more stable than PLPs made from WT coat protein, which denatured at 6 M urea (Figure 5B). However, PLPs made with the W410D coat variant were unstable, denaturing to coat monomers at 4 M urea. These data show that both I366 and W410 residues are involved in the stability of the PLPs.

The effect of substitutions at positions I366 and W410 on the PLP expansion was determined, as described above. PLPs with I366A substitution expanded at approximately 67 °C, similar to the WT PLPs (Figure 4C). PLPs assembled with I366D coat protein were unable to expand; at 72 °C; the PLPs simply rupture and run as an aggregate smear on the agarose gel. I366R PLPs are improperly assembled due to the substitution, resulting in a smeared band that runs higher than the WT PLP band. PLPs from the 22 °C samples, as well as the temperature at which they underwent expansion *in vitro*, were observed by electron microscopy (Figure 4D). All of the 22 °C samples were similar to the WT sample at this temperature, save the I366R sample that is composed mostly of aggregates. The samples at the expansion temperatures were all similar to WT, with the exception of I366R.

PLPs made with W410R coat protein expanded at 67 °C similar to PLPs made with WT coat protein (Figure 5C), whereas the W410A coat variants heat expanded at 63-67 °C. The morphology of the expanded W410R and W410D PLPs was altered compared to with WT or W410A PLPs, suggesting an effect on particle stability. W410D PLPs at 22 °C have some aberrant PLPs (green) in addition to the regularly formed structures (Figure 5D). The PLPs of the W410D coat variant reproducibly heat expanded incrementally, observed by the slight change in migration with each increase in the temperature, suggesting a decrease in the cooperativity in capsid expansion. PLPs with W410A substitutions in their coat proteins looked similar to PLPs assembled WT coat protein, both at 22 °C and at the expansion temperature. Thus, these data show that residue W410 plays a role in capsid maturation. These data also show that the hydrophobic pocket made of I366 and W410 is sensitive to charged substitutions, but not to a decrease in size of a non-polar residue.

#### Phages with I366 and W410 non-lethal coat substitutions are significantly less stable than those with WT coat protein

Since we observed significant effects of amino acid substitutions at W61, W410 and I366 on the stability of PLPs and the ability of PLPs to undergo expansion, we tested if phages assembled with the various non-lethal coat protein substitutions were also destabilized using a urea titration as described in the Methods. In this instance, only coat protein variants that are able make phages at some temperature could be assessed. The concentration of urea at which 50% of the initial titer remained after 16 hours of incubation was determined for phages assembled with WT coat protein or with W61Y, I366A, W410R, W410A, W410D coat substitutions (Figure 6, Table 2). Phages with WT coat protein declined to 50% of the initial titer at < 3 M urea. The W61Y and W410D phages were more stable than WT phages, with a 50% decline in titer at 3.5 M and 3.9 M urea, respectively. Conversely, phages with coat substitutions I366A, W410R, and W410A were all less stable than WT. Therefore, we show that the hydrophobic residues that interact with W61 within a capsomer greatly affect PLP and virion stability.

**Figure 6.**
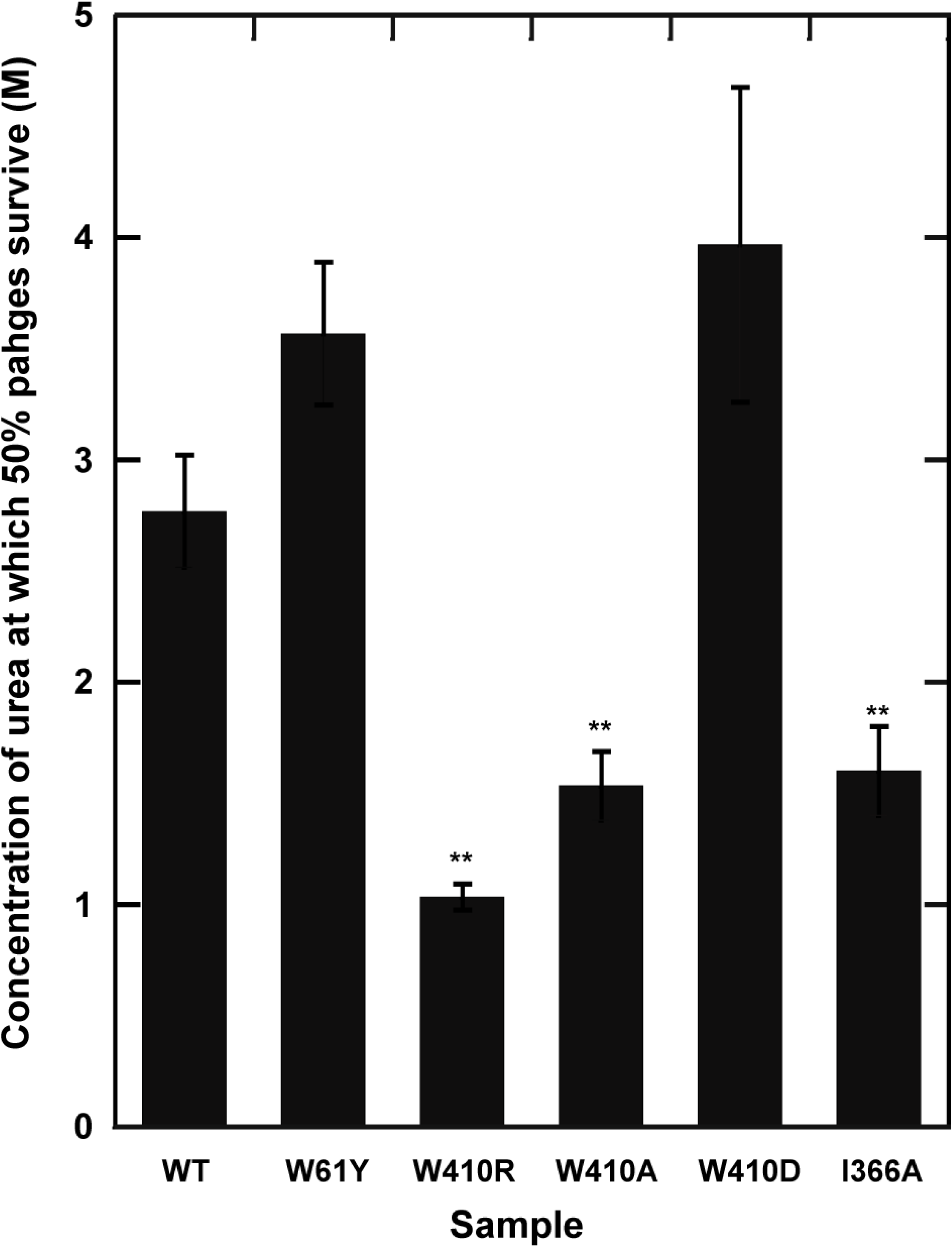
W410R and I366A substitutions affect virion stability. Bar graph depicts the urea concentration at which 50% of the phage population is remaining after 16 hours of incubation. These were calculated relative to the plaques formed at 0 M urea concentration. **A two-sample student t-test with unequal variance was done to calculate p<0.05.

### Interactions made between E-loop W61 with residues across the icosahedral two-fold axes of symmetry stabilize the capsid

Based on the cryoEM structure of the P22 capsid (30), W61 in the E-loop of one capsomer could be contacting residues A91, D92 and L401 in the P-domain loops of the neighboring capsomer (Figure 1). Distances between the residues were predicted using Chimera (38) and are all < 5 Å (Table 1). We hypothesize that the interactions between W61 of one capsomer and A91, D92 and L401 in a subunit of the adjacent capsomer could further seal the capsomers together, bolstering the stability of the capsid.

#### Formation phage particles is affected by substitutions at positions A91 and L401

Substitutions at positions A91, D92 and L401 were made in plasmid encoded gene 5 to test their effects on phage biogenesis using complementation of the *5-am* phage strain in the EOP assay, as described above (Figure 7A, Table 2). Mutating position D92 to ala or arg generated a WT phenotype. The A91V substitution yielded in a WT phenotype, while the A91D substitution resulted in a *cs* phenotype at 22°C. Complementation with the L401A coat substitution resulted in a *cs* and strong *ts* phenotype. The L401D coat protein substitution led to a lethal phenotype (Figure 7A). Thus, our data show that substitutions at A91 and L401 have the adverse effects on phage biogenesis.

**Figure 7.**
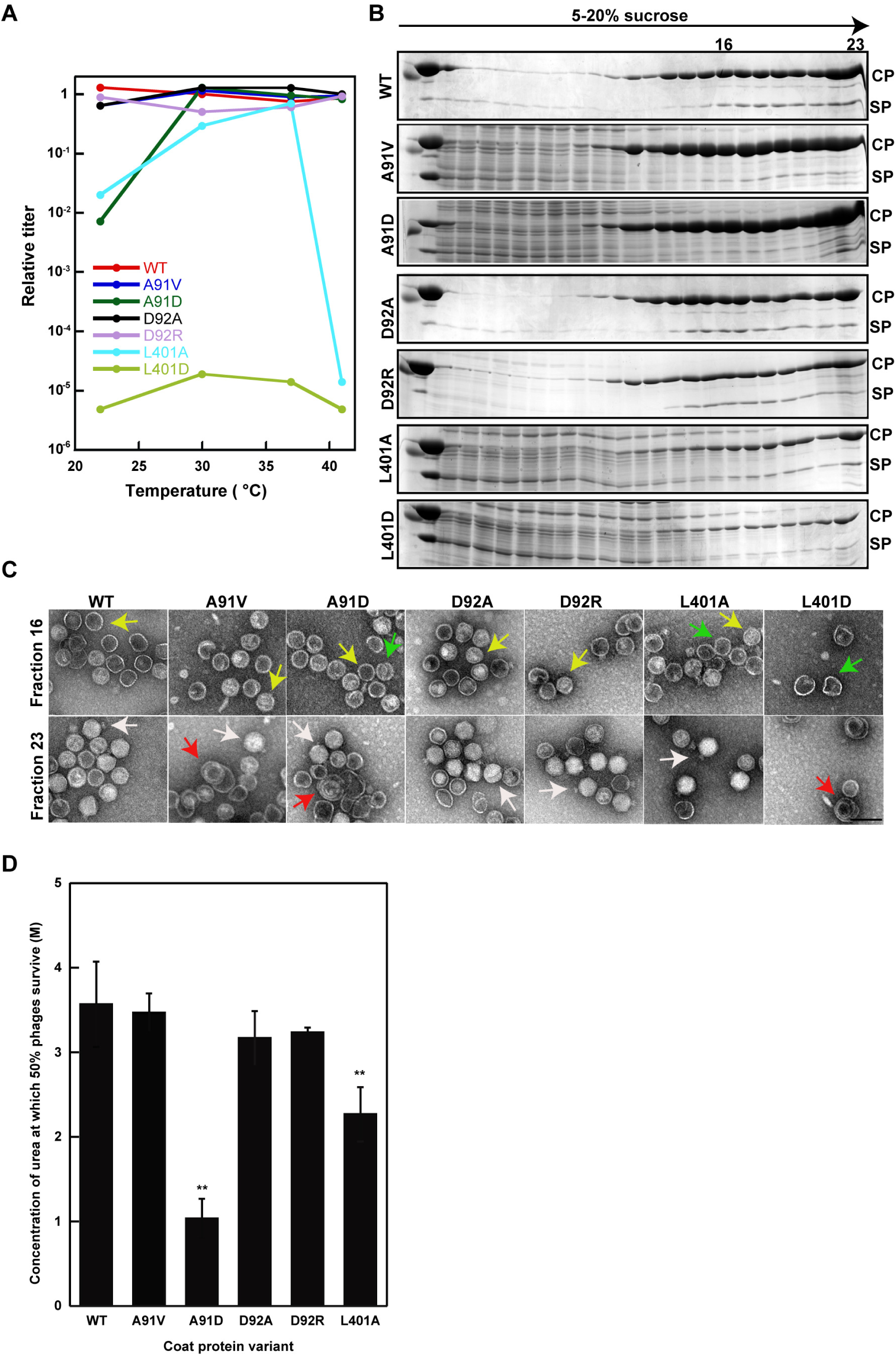
A91 and L401 affect procapsid assembly and capsid stability. **A.** Titers of phages assembled by complementation with A91V (blue), A91D(green), D92A (black), D92R (lavender), L401A (cyan) and L401D (light green) coat substitutions relative to phages with WT (red) coat protein at 30°C. **B.** SDS gels of 5-20% sucrose density gradient fractions from a phage-infected cell lysate to separate procapsids and phages with WT coat protein and coat protein with the A91, D92 and L401 coat substitutions. Fraction 16 is where normal procapsids sediment and fraction 23 is where phages and other particles are found. CP, coat protein; SP, scaffolding protein. **C. Top row**. TEM images of procapsid fractions (fraction 16) from sucrose density gradient of WT and A91, D92, L401 coat variants. **Bottom row**. Mature phage fractions (fraction 23) from the sucrose density gradient observed under the TEM. Yellow arrows show T=7 sized procapsids, green arrows show aberrant particles sedimenting at fraction 16, white arrows are highlighting mature phages and red arrows are highlighting aberrant particles sedimenting to the bottom of the gradient. Scale bar is 100 nm. **D.** Graph depicting the concentration of urea at which there is a 50% decline in the population of the samples. **Two-sample student t-test with unequal variance was used to calculate the p<0.05.

#### A91 and L401 coat protein substitutions result in the formation of aberrant particles

Procapsid and phage samples were made by complementation with coat protein having substitutions at positions A91, D92 and L401 from a lysate using a 5^-^13^-^*am* phage infection, as described in Methods. The samples were separated using a 5-20% sucrose gradient and fractions run on a 10% SDS gel. Procapsids sediment at approximately fraction 16 and phages sediment to the bottom (fraction 23) (Figure 7B). Fractions 16 and 23 were viewed by negative stain TEM (Figure 7C). All of the coat protein variants were all capable of assembling normal sized procapsids, as seen by both sucrose gradient and TEM (highlighted by yellow arrows in the micrographs), although the A91 and L401 variants also showed aberrant particles (green arrows). Only L401D coat protein was unable to support production of phages (phages indicated with white arrows), consistent with the lethal phenotype. These data suggest that there may be a hydrophobic interaction of A91 and L401 with W61 that affects assembly of procapsids.

Next, we tested if the non-lethal amino acid substitutions at these sites altered the stability of the viruses by treating them with different concentrations of urea, as described above (Figure 7D). Phages were isolated by complementation of a 5^-^13^-^*am* phage strain at 30 °C, as described in the Methods. Phages assembled with A91V, D92A and D92R coat protein variants had stabilities similar to WT coat protein phage samples, with all them having a 50% surviving population at approximately 3.5 M urea. However, virions with A91D and L401A coat protein substitutions showed a 50% drop in their populations at urea concentrations of 1 M and 2.2 M, respectively. Therefore, we show that residues A91 and L401 play an important role in stabilizing the capsid and likely does so by making intercapsomer contacts with W61.

## DISCUSSION

### The role of the E-loop in stabilizing capsids built with coat proteins having the HK97 fold

Capsid assembly and stability are governed by inter- and intra-capsomer interactions. In compliance with the local rule theory, the conformation of a subunit is dependent on the conformation of its neighboring subunits (39). Procapsid subunits additionally change conformation upon maturation, which also affects stabilizing interactions both within and between subunits. The burial of hydrophobic residues is an entropic process and is likely set into motion by capsid assembly, which is initially driven by weak interactions (40, 41). There is a greater percentage of hydrophobic amino acids at subunit interfaces, rather than the exterior of oligomeric proteins (42, 43). Indeed, P22 capsids are estimated to be stabilized by approximately – 24 kcal/mol subunit from intra-capsomer interactions due to buried hydrophobic residues along the edges of subunit and through E-loop interactions (44).

The E-loop has been implicated in making contacts with the P-domain and A-domain of the adjacent subunits in phages such as phi29 (20),T4 (10, 23), T7 (22), HSV-1 (21) and 80 alpha (45). It also makes intercapsomer contacts in the capsid of the HK97 phage that are important for procapsid assembly and are only found in procapsids (29). Intracapsomer contacts made by E-loops are important for proper conformation of capsomers in procapsids, and intercapsomer contacts are required for higher order assembly of capsomers to form a stable capsid (16). We show that a hydrophobic residue, W61, present in the phage P22 E-loop facilitates capsid stabilization. Since the E-loop of one subunit overlaps with the P-domain of the neighboring subunit within a capsomer, as well as reaches to the adjacent capsomer across two-fold axes of symmetry, we propose that W61 is involved in stabilizing the capsid, as described below. Table 2 provides a summary of all the mutants in this study, along with their respective phenotypes and mature virion capsid stabilities denoted by specific urea concentrations at which their populations decreased to half their original number.

### P22 capsids are stabilized by hydrophobic intracapsomer interactions

We propose that in phage P22, W61 in the E-loop may interact with residues I366 and W401 from the neighboring subunit within a capsomer, like a peg inserted into a hydrophobic pocket. The effects of switching the tryptophan at position 61 to asparagine or valine resulted in a lethal effect on the phage production. Furthermore, W61N and W61V PLPs do not expand *in vitro*, implicating this residue in maturation. When the W61 is replaced with tyrosine, the hydrophobic peg is maintained due to the size and the hydrophobicity of tyrosine and leads to a phenotype similar to WT. Coat protein monomers with these W61 substitutions were able to assemble, and interact with scaffolding protein, suggesting that the folding of the monomers was not the reason for any phenotypes. Substitutions at position 410 from tryptophan to arginine or aspartate resulted in a mild *cs* and *ts* phenotype, while W410A had a phenotype similar to WT, indicating that the size of the non-polar residue was not critical but that the replacement by a charged residue was not favored. PLPs generated W410D substitutions heat expanded at a temperature lower than WT showing that this residue affects capsid maturation. I366R and I366D coat substitutions cause a lethal phenotype. Phages assembled with W410R, and I366A coat protein mutants were less stable than phages with WT coat protein. Surrounding these hydrophobic interactions are several charged residues (Figure 1B), which we suggest further strengthens the interaction.

### Intercapsomer interactions that are crucial to P22 stability

Residue W61 also likely interacts with A91 and L401 of a subunit in the adjacent capsomer across two-fold axes of symmetry. Phages assembled with mutant coat proteins A91D, L401A had a *cs* and *ts* phenotype, while L401D resulted in a lethal phenotype. Conversely, D92A and D92R substitutions had a phenotype similar to WT, indicating that this residue is probably not involved in intercapsomer stabilization. The stability of phages assembled with coat protein mutants (in those cases where phages were produced) was tested by incubation in urea. Phages with A91D and L401A coat substitutions had a decreased capsid stability, suggesting that residues A91 and L401 likely participate in the association with W61. This was further confirmed by the formation of aberrant particles *in vivo* formed with A91D, A91V and L401A coat substitutions.

### Comparison of the stabilizing network between the P22-like phages

The importance of W61 in procapsid assembly and maturation is apparent when comparing coat proteins in the cluster of P22-like phages that includes 78 unique phages (46) (Figure S1). The coat protein sequences in the cluster vary widely with different branches having only 15-20% coat protein sequence identify. Residues at position 61 are conserved within phylogenetic branches and divergent between these branches. For example, within the P22-like phage cluster, the P22 major branch has tryptophan at position 61, Sf6 major branch has glycine and CUS-3 major branch has leucine at this position in the alignment (46). Similarly, residues at positions 91, 92, 366, 401 and 410 are also conserved within the major branches, but differ between the branches (Figure S1). The residues at these positions in each of these branches are shown in Table 3. Members in the CUS-3 major branch likely use a hydrophobic network to stabilize their capsids because they also possess hydrophobic residues at the positions that are involved in this interaction in the P22 capsid. The members of the Sf6 major branch either use electrostatic interactions to stabilize their capsids or use hydrophobic interactions in other regions of their coat proteins as a mechanism of capsid stability.

**Table 3.**
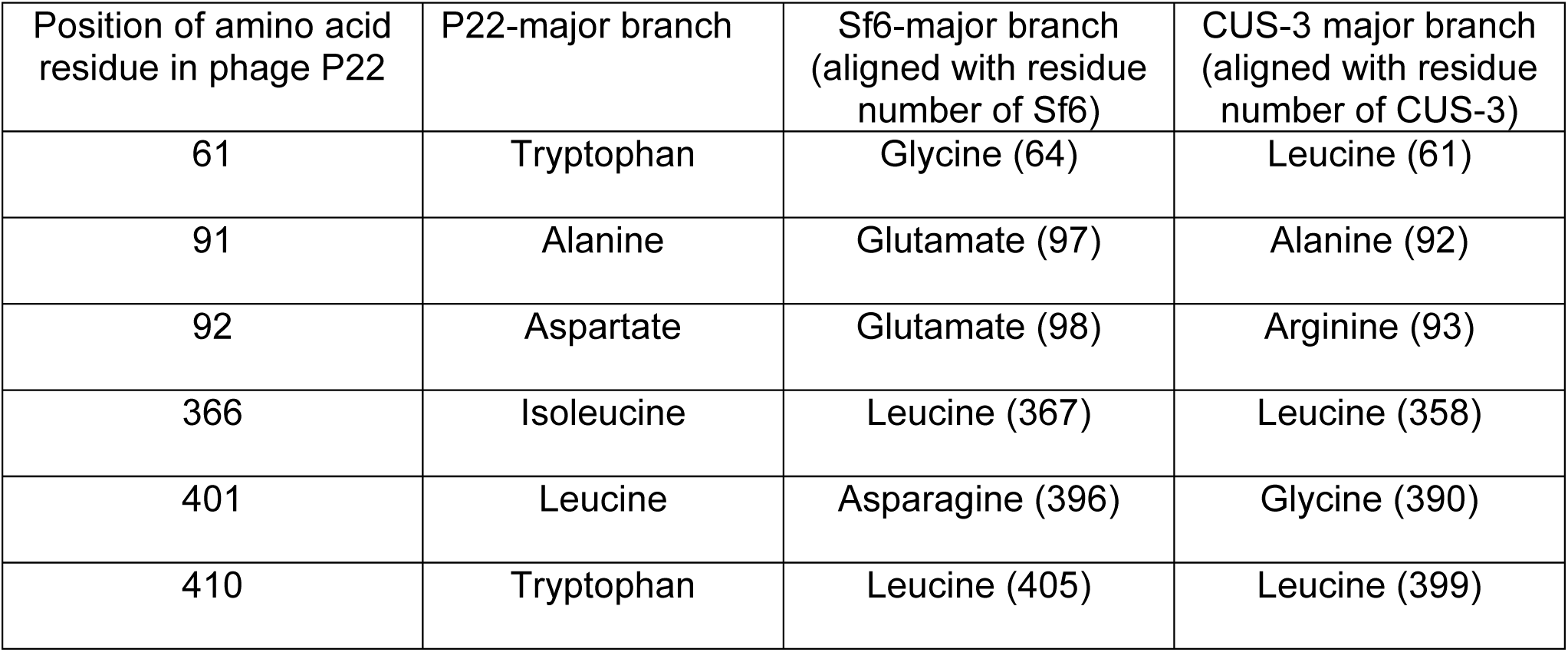
Comparison of hydrophobic peg forming residues in the P22 capsid with residues at those positions in the P22, Sf6 and CUS-3 major branches.

| Position of amino acid residue in phage P22 | P22-major branch | Sf6-major branch (aligned with residue number of Sf6) | CUS-3 major branch (aligned with residue number of CUS-3) |
| --- | --- | --- | --- |
| 61 | Tryptophan | Glycine (64) | Leucine (61) |
| 91 | Alanine | Glutamate (97) | Alanine (92) |
| 92 | Aspartate | Glutamate (98) | Arginine (93) |
| 366 | Isoleucine | Leucine (367) | Leucine (358) |
| 401 | Leucine | Asparagine (396) | Glycine (390) |
| 410 | Tryptophan | Leucine (405) | Leucine (399) |

In conclusion, our data suggest that phage P22, and members of the P22- and CUS-3 major branches of the P22-like phages, are stabilized by a hydrophobic network between the tip of the E-loop, which inserts into a hydrophobic pocket in the β-strands of the P-domain of a neighboring subunit within in a capsomer, and also with P-domain loops across two-fold axes of symmetry between capsomers. In total, these interactions within and between capsomers act like a hydrophobic net holding the capsid together. The interaction is further strengthened by a cage of charged residues surrounding the hydrophobic pocket. This mechanism is akin to the covalent crosslinking reaction that stabilize the HK97 capsid (15), where the site of crosslinking between polar residues K169 and N356 is protected by a hydrophobic cage composed of the amino acids leucine, methionine and valine (47). The CUS-3 major branch of the P22 like phages may use a similar method to stabilize capsids as P22 but how the Sf6-like branch of the P22-like phages has modified this network to produce stable capsids will require further investigation.

## MATERIALS AND METHODS

### Bacterial and phage strains

*Salmonella enterica* serovar Typhimurium strain DB7136 [*leuA414* (Am) *hisC525* (Am) *sup*^0^] (48) was used as the host for the EOP/complementation assay and for *in vivo* generation of procapsids and phages. *Salmonella enterica* serovar Typhimurium strain DB7155 was used for urea titration assays to test for capsid stability of mature virions. Strain DB7155 (*supE20 gln, leuA414*^*-*^*am, hisC525*^*-*^*am*) is a su^+^ derivative of DB7136 (48). The 5^-^*am* strain of P22 (5^-^*am* N114) contains an amber mutation in *gene 5* which codes for coat protein, while the *5*^*-*^*13*^*-*^*am* strain of P22 (5^-^*am* N114, 13^-^*am* H101) also has an amber mutation in the gene encoding the holin protein (gp13), which is responsible for cell lysis. All the P22 strains contain the allele c1-7, which ensures they do not enter lysogeny.

### Plasmids

Gene 5 was cloned between the BamHI and HindIII sites of plasmid pSE380 (Invitrogen) to form recombinant plasmid pMS11 (26). This plasmid was used for *in vivo* complementation. Plasmid pPC (derived from pET3a, kindly given to us by Dr. Peter E. Prevelige) has gene 8 (encoding scaffolding protein) followed by gene 5 (49). Site-directed mutagenesis was used to generate mutations in the gene 5 of both the plasmids.

### Efficiency of plating (EOP) complementation assay

Strain DB7136 containing pMS11 with the substitution of interest was plated with 5^-^*am* phage. The *Salmonella* cells were grown to mid-log phase, harvested by centrifugation and concentrated by resuspending in a small volume of LB. The cells were then mixed with 5^-^*am* phage in soft agar containing 1 mM IPTG and poured over LB plates containing ampicillin. The plates were incubated at 22 °C, 30 °C, 37 °C and 41 °C. The relative titer of phages with coat substitutions was calculated by counting their plaques at the temperature being tested, relative to the 5^*-*^*am* phages complemented with WT coat protein expressed from the pMS11 at permissive temperature (30 °C).

### *In vivo* generation of procapsids and virions

*Salmonella* strain DB7136 containing pMS11 with the coat substitutions in gene 5 was grown until mid-log phase (∼2 × 10^8^ cells/ml). The cells were then infected with 5^-^*am* 13^-^*am* phage at a multiplicity of infection (MOI) of 5. The coat protein expression was simultaneously induced with 1 mM IPTG. The cells were grown for an additional four hours, harvested and stored overnight at −20 °C after being resuspended in a lysis buffer (50mM EDTA, 0.1% Triton X-100, 200 μg/ml lysozyme, 50 mM Trizma base, 25 mM NaCl, pH 7.6). The cells were processed as described previously (26). Briefly, the cells underwent two cycles of freezing and thawing at room temperature. DNase and RNase were added at 100 μg/ml and phenylmethylsulfonyl fluoride (PMSF) was added at a final concentration of 1 mM. The supernatant retrieved from separating out the debris was then centrifuged at 17901 X g in RP80-AT2 rotor (Sorvall) for 20 min to pellet the procapsids and phages. The pellet was then suspended overnight in 20 mM sodium phosphate buffer (pH 7.6) containing 20 mM MgCl_2_.

### Preparation of procapsid–like particles

Procapsid-like particles (PLPs) are assembled from coat protein and scaffolding protein expressed from pPC, which has gene 8 followed by gene 5 in pET3a (49). The plasmid-containing BL21 (DE3) cells were grown in LB in ampicillin at a final concentration of 100μg/ml. Expression of T7 polymerase was induced by the addition of 1 mM IPTG, and the cells grown for an additional 4 hours. The PLPs were pelleted from the lysed cells and purified over a sizing column containing Sephacryl S1000 matrix (GE Healthcare).

### Preparation of coat protein monomers

To make coat monomers, the scaffolding protein is first extracted from the PLPs, resulting in empty coat protein shells, as described previously (49, 50). To generate monomers, the empty shells were unfolded in 6.75 M urea, in 20 mM phosphate buffer pH 7.6. The denatured protein was dialyzed in Spectra/Por dialysis tubing (Molecular weight cutoff: 12-14 kDa) two times for three hours and once overnight against 20 mM phosphate buffer, pH 7.6 at 4°C. The refolded coat protein monomers were centrifuged in a RP80-AT rotor (Sorvall) at 4 °C at 60,000 rpm for 20 minutes to remove aggregates or assembled particles. The supernatant contained the active coat monomers.

### Sucrose Density Gradients

A 5-20% sucrose density gradient was prepared by using a gradient maker (model 106; Biocomp Instruments). A 100μl of sample containing a mixture of procapsids and virions was applied to the top of the gradient, and centrifuged in the RP55S rotor of the RC-M120 EX centrifuge (Sorvall) at 104813 X g for 35 minutes at 20 °C. The gradients were fractionated by hand from the top into 23 100 μl fractions.

### Cesium chloride gradients

Cesium chloride gradients were made to separate the procapsid and mature virus mixtures that were generated *in vivo* as described above. 25% sucrose, 1.4 gm/cc and 1.6 gm/cc cesium chloride solutions were prepared in 20 mM phosphate buffer (pH 7.6) to prepare the gradient. The gradient was made by successively layering 25% sucrose solution on top of a 1.4 gm/cc cesium chloride solution, which was further layered on top of a 1.6 gm/cc cesium chloride solution. The procapsid and phage mixture was applied to the top of the layered gradient and centrifuged in Sorvall MX 120 ultracentrifuge for an hour at 30,000 rpm at 18°C. Phages sediment to the interface of the 1.4 gm/cc and the 1.6gm/cc layers of cesium chloride. Phages were extracted using a syringe and used for the urea titration assay to test for capsid stability.

### Weak affinity chromatography

Hexa-histidine-tagged scaffolding protein (6 mg) was loaded on a 1 ml immobilized metal affinity chromatography column charged with cobalt (Clontech). Coat protein monomers (0.2 mg/ml) were loaded on to the column, run at a flow rate of 1.25 ml/min and 0.25 ml fractions were collected. Ovalbumin was used as the negative control for non-specific binding and was loaded at 0.2 mg/ml. Tryptophan fluorescence of the collected fractions was then measured using Horiba Fluoromax 4 fluorometer with the excitation wavelength at 295 nm and a bandpass of 1 nm and emission wavelength set at 340 nm with a bandpass of 8 nm. The fluorescence readings were recorded was arbitrary units (A.U.).

### Circular dichroism spectroscopy

Spectra of prepared coat protein monomers were observed using the Applied Photophysics Pistar 180 spectrapolarimeter (Leatherhead, Surrey, United Kingdom) using a cuvette with a 1 mm path length. The spectra were taken at a concentration of 0.3 mg/ml in 20 mM phosphate buffer (pH 7.6) at 20 °C. Scans were done with a1 nm step size between wavelengths 195 nm and 225 nm. The bandpass was 3 nm and the time-per-point averaging was set to 55 sec. The molar ellipticity was used to compare secondary structure of coat variants with that of WT coat monomers.

### Negative stain microscopy

3 μl of the samples from fractions 16 and 23 of the sucrose gradient were spotted onto carbon-coated copper grids (Electron Microscopy Sciences). They were then washed with 2-3 drops of water, stained with 1% uranyl acetate 30 seconds and the excess blotted. A Tecnai Biotwin transmission electron microscope was used to observe the grids at 68000X magnification.

### Urea titration of PLPs

9 M urea prepared in phosphate buffer at pH 7.6 was used to make 1-7 M urea at 1 M intervals. The refractive index of the prepared urea was checked using a refractometer to confirm the concentration. PLPs were then diluted in each concentration of urea to a final concentration of 0.5 mg/ml. The samples were left overnight in urea and about 5 μg of the sample was run on a 1% agarose gel using SeaKem LE Agarose in 1X Tris-acetate-ethylenediaminetetraacetic acid (TAE) buffer.

### Heat expansion

PLPs (1 mg/ml) were incubated at temperatures ranging from 25 °C to 72 °C for 15 min and then placed on ice. The samples were then run on a 1% agarose gel in 1X TAE buffer.

### Urea titration of mature virions to test for capsid stability

Phages generated *in vivo* and purified using cesium chloride gradient were used for this assay. After the determination of the titer of these phages, 10^4^ phages/ml were treated with urea concentrations from 0-8 M. A 9 M stock solution of urea, prepared in 20 mM phosphate buffer, was used to make the appropriate dilutions for the assay. The phage and urea mixture were incubated overnight at 22°C following which 10μl of the incubated sample was mixed with 3 drops of DB7155 strain of *Salmonella enterica* in 2.5ml of soft agar. The solution was poured over plain LB plates and incubated overnight at 30°C. Plaques were enumerated after overnight incubation.

## ACKNOWLEDGMENTS

This research was supported by NIH grant R01 GM076661. We would also like to thank Dr. Xuanhao Sun and Maritza Abril from the electron microscopy facility at the University of Connecticut for their assistance with the electron microscope and Dr. Nikhil Ram Mohan (Boston College) for help with the sequence analyses of the P22-like phage cluster.

